# Transcriptomic cell type structures in vivo neuronal activity across multiple time scales

**DOI:** 10.1101/2022.07.10.499487

**Authors:** Aidan Schneider, Mehdi Azabou, Louis McDougall-Vigier, David Parks, Sahara Ensley, Kiran Bhaskaran-Nair, Tom Nowakowski, Eva L. Dyer, Keith B. Hengen

**Author notes:** Contributed equally as co-first authors (Aidan Schneider, Mehdi Azabou). Correspondence should be sent to: Keith Hengen and Eva Dyer.

## Abstract

**SUMMARY:** Cell type is hypothesized to be a key determinant of the role of a neuron within a circuit. However, it is unknown whether a neuron’s transcriptomic type influences the timing of its activity in the intact brain. In other words, can transcriptomic cell type be extracted from the time series of a neuron’s activity? To address this question, we developed a new deep learning architecture that learns features of interevent intervals across multiple timescales (milliseconds to >30 min). We show that transcriptomic cell class information is robustly embedded in the timing of single neuron activity recorded in the intact brain of behaving animals (calcium imaging and extracellular electrophysiology), as well as in a bio-realistic model of visual cortex. In contrast, we were unable to reliably extract cell identity from summary measures of rate, variance, and interevent interval statistics. We applied our analyses to the question of whether transcriptomic subtypes of excitatory neurons represent functionally distinct classes. In the calcium imaging dataset, which contains a diverse set of excitatory Cre lines, we found that a subset of excitatory cell types are computationally distinguishable based upon their Cre lines, and that excitatory types can be classified with higher accuracy when considering their cortical layer and projection class. Here we address the fundamental question of whether a neuron, within a complex cortical network, embeds a fingerprint of its transcriptomic identity into its activity. Our results reveal robust computational fingerprints for transcriptomic types and classes across diverse contexts, defined over multiple timescales.

## INTRODUCTION

Mammalian brains are composed of between 10^8^ to 10^11^ neurons that can be categorized into hundreds of genetically distinct cell classes and types (Scala et al., 2021; BRAIN Initiative Cell Census Network, 2021). There is great interest in understanding the role of cell types in the intact brain, driven by the assumption that a careful description of cells, their anatomical diversity, and their rules of connectivity will offer insight into how circuits generate behavior (Ecker et al., 2017; Brain Initiative Cell Census Network, 2021; Yuste et al., 2020). Anatomically and morphologically diverse classes of neurons were initially described by Ramon y Cajal (1892), while more recently, neuronal classes have been defined based on molecular markers, morphology, and anatomical location. Single cell transcriptomics has revealed a diverse but finite number of profiles that align with known cell classes, thus indicating the genetic basis of cellular phenotype (Lein et al., 2017; Tasic et al., 2016; BRAIN Initiative Cell Census Network, 2021). How diverse cell types influence the pattern and timing of neuronal spiking in the intact circuit is largely unknown (Lein et al., 2017).

Previous attempts to determine the functional signature of transcriptomic cell types have taken advantage of reduced preparations and synaptically isolated cells. Ex vivo patch clamp-based studies suggest that, given a controlled input, electrical properties of neurons can also provide insight into genetic identity (Nowak et al., 2003; Rainnie et al., 2006; Gouwens et al., 2020; Lazarevich et al., 2021). However, finding reliable signatures of cell types in the intact brain has proven elusive because isocortical spike timing is highly irregular (Softky and Koch, 1993; Mainen and Sejnowski, 1995; Stevens and Zador, 1998; Nolte et al., 2019; Seeman et al., 2018; Peron et al., 2020) and even distinct neuronal classes exhibit broad variability in their activity patterns (Bartho et al., 2004; Cardin et al., 2007; Royer et al., 2012; Buzsáki and Mizuseki, 2014). The challenge of identifying spike-timing-based signatures of neuronal classes is exacerbated by experimental limitations related to simultaneously measuring physiology and genetic cell type (Lima et al., 2009; Ding et al., 2022). As a result, classification of cell type in the intact brain relies either on a) molecular tools or b) analysis of the shape of the extracellular waveform, which indicates only parvalbumin (PV) positive neurons amongst additional clusters that are currently not tied to a transcriptomic cell type (Bartho et al., 2004; Niell and Stryker, 2008; Jia et al., 2019; Trainito et al., 2019).

Ongoing developments in genetic tools and high density recordings are accelerating the investigation of how cell types impact brain dynamics. Recent work demonstrates that inhibitory transcriptomic cell type influences activity correlations amongst subpopulations of neurons (Bugeon et al., 2022). This work reveals that molecularly diverse cell classes result in similarly diverse patterns of population activity. This powerful observation underscores an important question: if cell type is observable in population interactions, does it impose reliable operational constraints at the level of individual neuron activity? In other words, what is the minimal unit at which transcriptomic class impacts cortical computation? Should a spike train from an individual neuron carry identity information, cell type would be discernible by any observer of spiking, such as an experimenter or the cortex itself. In this case, cell type information would only require an individual time series without reference to the time-locked, context-dependent activity of multiple populations.

Neuronal classes are typically described in terms of *global features*, such as firing rates (Niell and Stryker, 2008; Liu et al., 2009; Hengen et al., 2013), or *local features* that arise in response to acute events, such as sharp wave ripples (Stark et al., 2014) or response to a stimulus (Runyan et al., 2010). We reasoned that computational fingerprints of genetic cell types might exist when features of neuronal activity can be learned across time scales. To investigate this possibility, we developed a novel deep learning architecture that employs an attentional mechanism to learn signatures of transcriptomic cell type from the time series of a single neuron’s activity when examined over multiple time scales (millisecond to >30 minutes) and across presentations of diverse stimuli. This approach accurately decodes 16 cell types in a bio-realistic realistic model of primary visual cortex with 250,000 neurons. In both electrophysiology and optophysiology datasets where transcriptomic types can be identified (or inferred), we show that the transcriptomic type can be decoded from the timing of neuronal activity in the isocortex of behaving animals. We then apply this tool to the question of whether genetic subtypes of excitatory neurons are computationally discernable in the intact brain. Specifically, we identified 1) robust spike-timing signatures of the three transcriptomic families of inhibitory neuron labeled in high density electrophysiological recordings (parvalbumin, somatostatin, and vasoactive intestinal polypeptide expressing), 2) robust event-timing signatures of the same three interneuron classes and the excitatory neuron class in calcium imaging experiments, and 3) robust event-timing signatures of a subset of excitatory subtypes in calcium imaging experiments. Finally, we demonstrate that the computational fingerprints of cell types may be persistent across diverse stimulus sets. By revealing that a neurons’ transcriptomic identity is embedded across multiple timescales of its activity, we open the possibility that this signal may be accessible to and implicated in computations throughout the broader cortical network.

## RESULTS

To examine whether information about transcriptomic cell type is embedded in the timing of isocortical neuronal activity in the intact circuit, we sought datasets that sampled the activity of a large number of neurons and contained ground-truth information about each neuron’s underlying transcriptomic type. These requirements were satisfied by two open datasets centered on mouse primary visual cortex from the Allen Institute, specifically high-density extracellular electrophysiology with optotagging of inhibitory types (Siegle et al., 2021) and optophysiology/Ca^2+^ imaging (de Vries et al., 2020). In addition, we deployed a bio-realistic model of primary visual cortex comprised of 230,000 neurons (50,000 in the model core) with 17 cell-types modeled on transcriptomic classes in a layer-dependent fashion (Billeh et al., 2020). In all of these cases, recordings were obtained from visual cortical regions under similar sets of tasks and conditions (Figure 1A-C). We refer to broad transcriptomic families (parvalbumin (PV), vasoactive intestinal peptide (VIP), somatostatin (SST) and excitatory (E)) as “cell class”, and more specific transcriptomic subdivisions of E neurons as “cell type”. Here we use the timestamps of spikes and detected calcium events from all neurons paired with information about the underlying transcriptomic identity. In the electrophysiological dataset, three inhibitory neuron classes were genetically identified with Cre driver lines as expressing either PV, VIP, or SST. In the calcium imaging dataset, in addition to the same three inhibitory classes, eight different E Cre lines were recorded across cortical layers (Cux2-L2/3, Ntsr1-L6, Rorb-L4, Fezf-L5, Nr5a-L4, Rbp4-L5, Scnn1a-L4, Tlx3-L5). All E Cre lines were recorded at a single imaging depth, with exception of Cux2 (recorded are both in L1, L2/3) (de Vries et al., 2020). In contrast to the inhibitory cell classes, excitatory Cre lines varied in their degree of underlying specificity, with some lines comprising sparse labels of a single cell type and others targeting multiple cell types and layers simultaneously. In the bio-realistic model, the same three inhibitory classes and an E class were specified by layer, each with functional and connectomic properties informed by experimental evidence (Billeh et al., 2020). In all of our experiments, neurons were divided into non-overlapping train, validation, and test pools. Class balancing was performed such that models did not learn to simply predict the most prevalent cell classes (see methods). As a result, chance is always 1/number of cell types (K). When appropriate, we report balanced accuracy, defined per class as: (true positive rate + true negative rate) / 2, which is well suited to assess model performance when the representation of classes is imbalanced (Brodersen et al., 2010).

**FIGURE 1:**
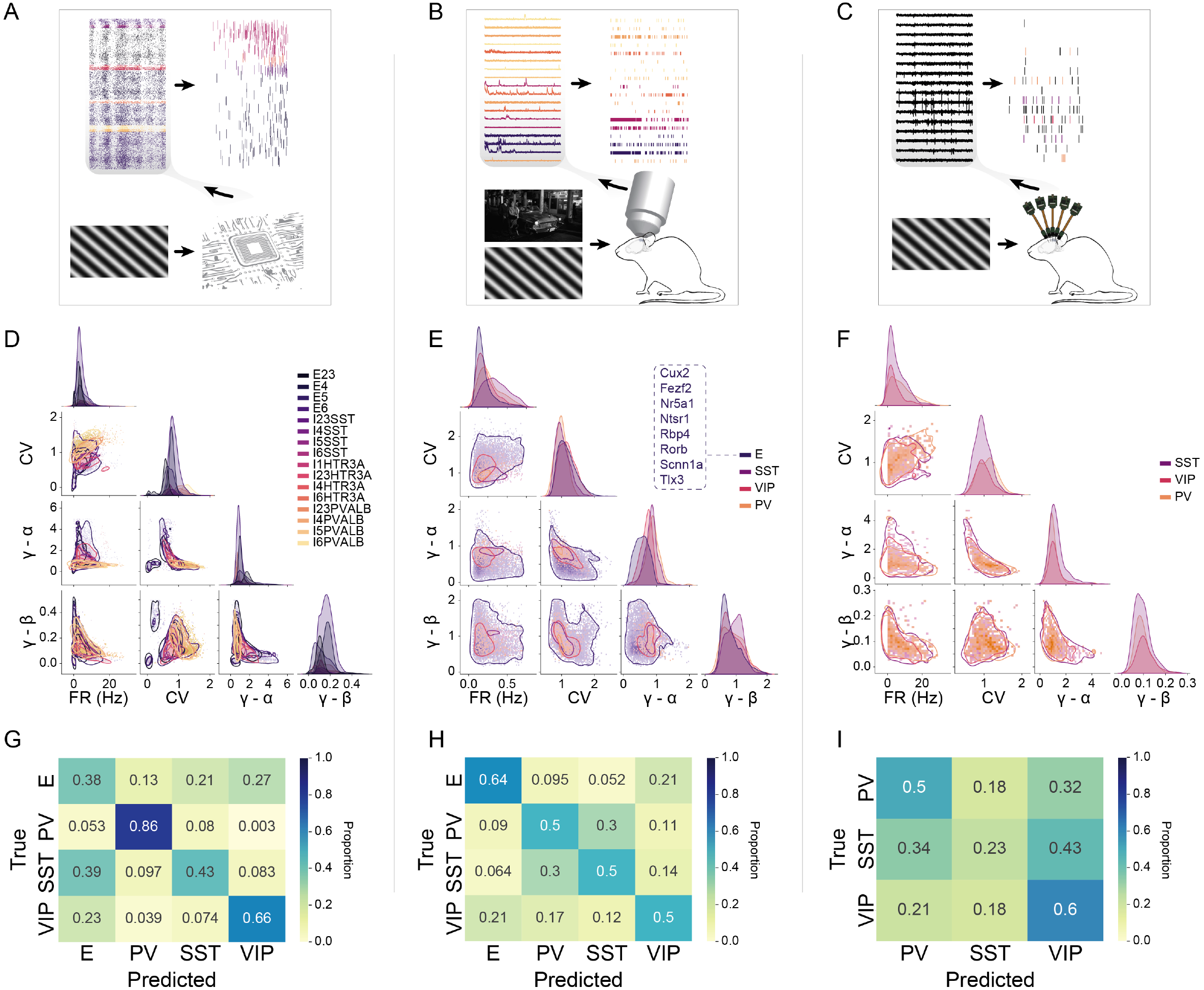
Experimental overview and linear separability of cell types. Three intact-circuit and/or in vivo datasets contain neuronal event time series with a corresponding genetic type. (A-C) An example visual stimulus, recording set-up, raw data trace, and extracted event raster are shown for: A) bio-realistic model of V1 (4 cell classes organized by layer to generate 16 different putative cell types), B) in vivo calcium imaging sessions acquired across three inhibi tory classes and eight excitatory cell types and C) extracellular electrophysiology from high-density Neuropixels containing three optotagged inhibitory classes (VIP, SST, PV). (D-F) Linear separability of cell type based on firing rate (FR) and three parameters extracted from the inter-event interval (IEI) distribution (coefficient of variation (CV), two gamma distribution parameters (a,) fit to the IEI distribution). (G-1) Confusion matrices for predicting cell class based on the combination of FR, CV, and gamma parameters.

### Cell types in summary statistics

Taking advantage of the extensive recordings of genetically labeled cells, we examined the degree to which cell type can be decoded from summary statistics of spiking (e.g., mean firing rate; FR, coefficient of variation (CV)) (Figure 1D-F). In each recording modality, we used a simple linear model (logistic regression) to empirically determine the maximum separability of cell type based on all combinations of the following summary statistics to examine functional differences in vivo: mean firing rate (FR), coefficient of variation (CV), and the two intrinsic parameters of the gamma distribution of interevent intervals (IEIs) (Li et al., 2018). In the V1 model, when we considered 16 cell types (K=16. Layer 5 Htr3a (VIP) had relatively too few neurons for inclusion; chance = 6.25%), FR, CV, and the gamma distribution parameters (ɑ, β) individually provided resolution slightly above chance (FR: 9.5 +/-0.3%, CV: 17.4 +/-0.5%, and ɑ, β: 25.9 +/-0.4%). In combination, these parameters resulted in a balanced accuracy of 31.5 +/-4.5%. For benchmark comparison to in vivo datasets, we collapsed the 16 cell types into K=4 (PV, VIP, SST, and E; chance = 25%). In this case, the combination of the three summary statistics yielded a balanced accuracy of 58.6 +/-0.2%. Similarly, in the electrophysiological dataset, the same parameters individually identified cell class (K = 3; chance = 33%) at or above chance (FR: 40.4 +/-0.9%, CV: 39.9 +/-0.7%, and ɑ, β: 34.0 +/-0.7%), and at 39.2 +/-1.1% balanced accuracy in combination. In the calcium imaging dataset, we collapsed the 10 excitatory cell types into a single class to consider the same level of prediction with K=4 (PV, VIP, SST and E; chance = 25%). FR, CV, and the gamma parameters individually identified cell class poorly (FR 35.3 +/-1.0%, CV: 31.3 +/-1.0%, ɑ, β: 43.1 +/-1.1%), and in combination achieved a balanced accuracy of 53.5 +/-0.9%. These results suggest that genetic cell type is loosely imprinted on global features of spike/event times, but that these metrics are insufficient to reliably infer the cell type of individual neurons.

### Nonlinear separation of cell types in IEIs

Summary statistics, such as CV, reduce many observed events to a single feature. We reasoned that consideration of the entire distribution of IEIs would either contain more information about genetic class, or prove to be too noisy to discriminate class beyond the basic descriptors. To distinguish between these possibilities, we trained a multilayer perceptron (MLP; see methods) to classify cell type/class based on a histogram of all IEIs generated by individual neurons. The MLP has the added advantage of not assuming linear separability of classes. In each dataset, we divided all labeled neurons into three groups split across individuals; 60% were used for training, 20% for validation, and 20% for testing. To account for results driven by sampling error, we trained and tested on 20 independent random splits of each dataset. In all contexts, the MLP-based approach vastly outperformed linear classifications based on summary statistics. In the full V1 model (K = 16), the MLP achieved 64.98% balanced accuracy (+/-0.26%), a 106.5% improvement over linear models (i.e. more than twice as accurate) (p=3.64E-40, ANOVA). Collapsed to four cell classes (E, PV, SST, and VIP), the MLP learned classification at a level of 80.76% (+/-0.28%), an improvement of 38.0% (p=1.85E-39, ANOVA). Similar improvements were observed in vivo. When trained on the K = 4 cell classes in the context of calcium imaging, the MLP achieved a balanced accuracy of 73.01+/-0.54%, an improvement of 36.4% (p=3.00E-16, ANOVA). In the electrophysiological dataset (K=3), the MLP learned to identify the three labeled inhibitory classes (PV, SST, VIP) at 49.52% balanced accuracy (+/-0.87%), an improvement of 10.6% (2.78E-4, ANOVA). The significant improvements observed in the MLP-based approach are driven by the computational advantage of being able to learn nuanced, nonlinear separation of classes in the full IEI distribution. However, this approach lacks any sequential information, such as patterns in IEIs or changes over time.

### LOLCAT

We hypothesized that it may be possible to learn cell type-specific fingerprints by considering the activity of a neuron at many time scales, both in terms of local event times and patterns in firing over longer time periods and across different stimuli. With this in mind, we developed a model (LOLCAT; **LO**cal **L**atent **C**oncatenated **AT**tention) that can both extract features from local snapshots of the event timing (brief IEI distributions) and learn global features of different cell types that may be resolvable over much longer timescales. LOLCAT learns to extract local features from short chunks of time before passing them through a multi-head attentional network that emphasizes certain trials through different “attention weights” and aggregates the weighted instance-level features into a final, global estimate of the cell type (Fig. 2A). The addition of multiple attention heads allows the model to extract features of activity at distinct yet complementary timescales from a single IEI to the extent of the recording.

**FIGURE 2:**
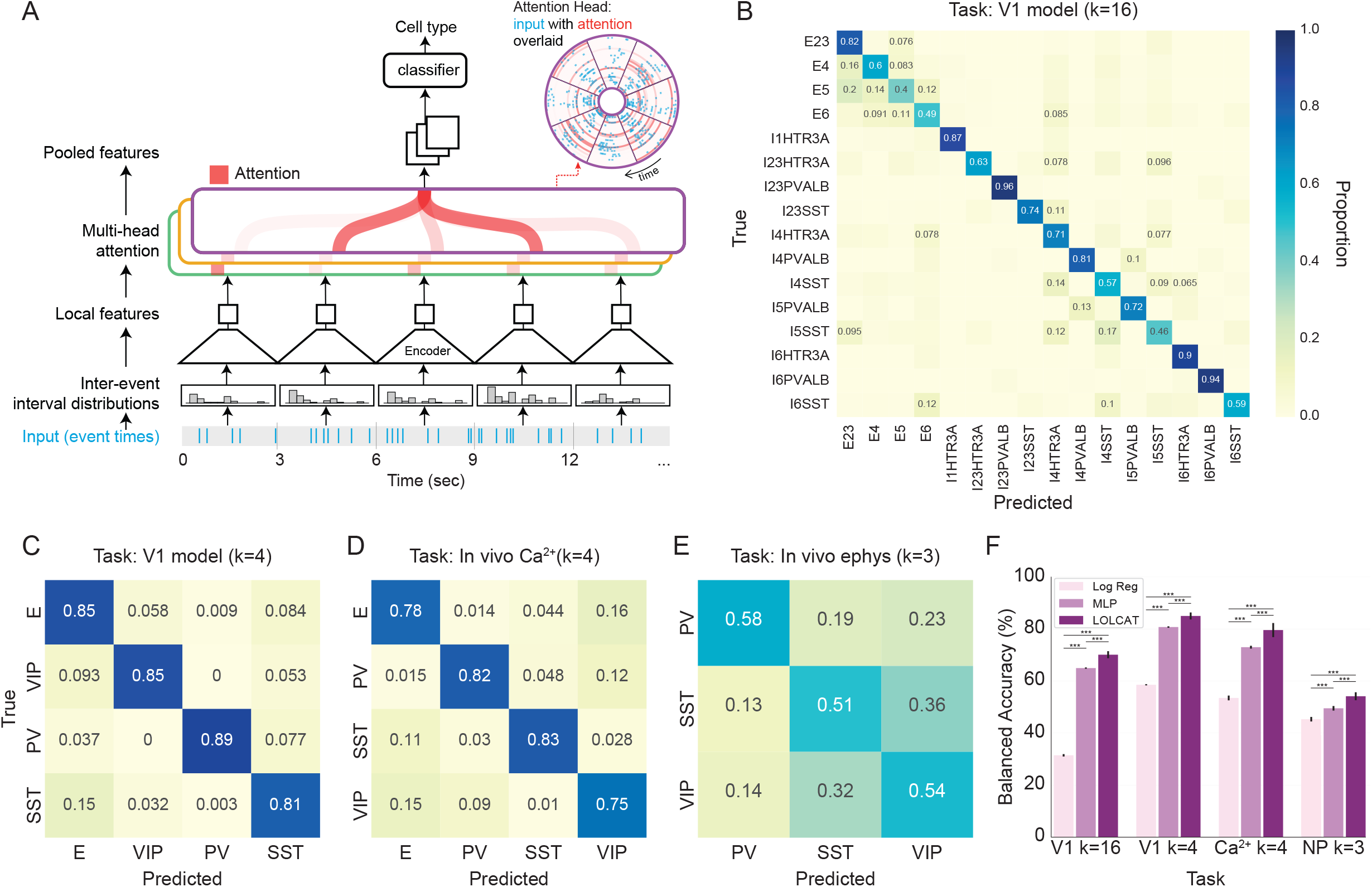
LOLCAT learns robust computational fingerprints of cell types. A) Architecture for LOLCAT: a time series of neuronal activity is split into short (3 sec) windows and the IEI distribution is estimated for each local window. IEIs are fed into an encoder to obtain features for each window before being aggregated in a multi-head attention module. The Attention Module consists of multiple “heads”, each of which computes a weighted combination of the representations of each IEI distribution, creating multiple pooled versions of the representations that are concatenated to obtain an estimate of the class. The network is trained in a supervised manner using dropout, both across the input blocks and at units in layers within the network. B) Confusion matrix for LOLCAT for the V1 model with 16 cell types, organized into 4 excitatory, and 12 inhibitory types (HTR3A is analogous to VIP). Annotation of performance at below chance is deleted for visual clarity. C,D) LOLCAT confusion matrix for identifying four broad cell classes, PV, VIP, SST and excitatory in the V1 model and in vivo calcium imaging data. E) LOLCAT confusion matrix for identifying three inhibitory cell classes, PV, VIP, and SST, in in vivo extracellular electrophysiological recordings. F) Summary of final balanced accuracy across different methods of decoding cell types from event times. Logistic regression (log reg) was trained on global statistics and IEI features. A multilayer perceptron (MLP; nonlinear deep network) was trained on the full IEI distribution. LOLCAT was trained on collections of IEI distributions observed over different windows and stimuli.

Starting with the in silico model, we first tested the ability of LOLCAT to learn fingerprints of all 16 cell types (K = 16)(Fig. 2B). In this context, LOLCAT achieved a mean balanced accuracy of 70.1 +/ - 1.4% (chance = 6.25%), a 7.9% improvement over the MLP (p=3.70E-15, ANOVA), and 122.9% over the linear model (p=3.64E-40, ANOVA) (Fig. 2F). The accuracy of individual cell types ranged between 40 and 96%. Out of 16 cell types, only five performed under 60%, and seven under 70% accuracy. When the 16 cell types were collapsed into the four genetic classes that most overlap with the in vivo datasets (K=4; PV, VIP, SST and E; chance = 25%), LOLCAT achieved a mean balanced accuracy of 85.0 +/-1.34%, with an even diagonal structure of the confusion matrix with values (PV: 89%, VIP: 85%, SST 81% and E 85%). This is a 5.3% improvement over the MLP (p=2.36E-12, ANOVA) and 45.1% over the linear model (p=1.61E-41, ANOVA).

Surprisingly, LOLCAT achieved a similar balanced accuracy, 79.59 +/-2.7%, trained on event timestamps from the calcium imaging data with the K=4 label set (PV, VIP, SST, E; chance = 25%)(Fig. 2C). This represents a 9.0% improvement of the MLP (p=1.04E-9, ANOVA) and a 48.8% improvement over the linear model (p=7.91E-20 ANOVA) (Fig. 2F). Model accuracy was well-balanced across classes (E: 78%, PV 82%, SST 83%, VIP 75%). The accuracy achieved for each class is noteworthy given the massive class imbalance between excitatory and scarce inhibitory cells in this dataset (PV: 180, SST: 470, VIP: 470, E: 22,245).

While the relationship between deconvolved event times in calcium recordings and action potentials is not one to one, this relationship may be preserved in electrophysiological recordings. Trained on spike times from optotagged neurons in the electrophysiological dataset (K=3; PV, SST, and VIP; chance = 33%), LOLCAT extracted genetic cell class with a balanced accuracy of 54.2 +/-1.5% (Fig. 2E). These results are a 9.4% improvement relative to the MLP (p=2.29E-4 ANOVA), and a 20.1% relative to the global statistics baseline (p=1.0 ANOVA)(Fig. 2F). Class accuracies were reasonably well-balanced (PV: 58%, VIP: 54%, and SST: 51%). Together, these results suggest that in the intact isocortical circuit, information about genetic cell type is robustly embedded in the time series of neuronal activity. It is worth noting that, for a variety of possible reasons, it is more challenging to extract transcriptomic class from this dataset when compared with the calcium imaging data.

Across all three datasets, LOLCAT resulted in well-balanced class identification (Fig. 2B-E). In the benchmark comparisons (in silico K=4; calcium imaging K = 4; electrophysiology K = 3) LOLCAT exhibited a range of class accuracies (highest class accuracy - lowest class accuracy) of 8, 8, and 7%, respectively. These results indicate that LOLCAT achieved increased accuracy by enhancing the resolvability of all classes rather than a subset. In contrast, accuracy ranges in the MLP results were 28, 32, and 36%. In the linear models, ranges of 47, 37, and 14% were observed.

### LOLCAT interpretability

Our model can extract both local features from spike and event intervals, as well as increasingly global activity patterns that emerge over longer timescales. Thus, we sought to understand the rules the model might be using to solve this cell type discrimination task. To understand how the model incorporates global information about neural activity, we first quantified the relationship between model confidence and the number of trials observed as a function of cell class in the calcium imaging dataset (Fig. 3A). Summarily, SST interneurons required relatively few observations, and VIP interneurons required the most. All four classes were significantly different from one another in this respect (p<0.001, Linear Mixed Model with Tukey posthoc). To understand the global features extracted from the network when neurons are studied over long timescales, we visualized the trial-based allocation of attention (i.e., attention masks) in the four attention heads in the model (Fig. 3B). This revealed that identification of classes was distributed across all four heads, and that some heads learned sparse attentional rules while others learned global rules. In other words, some heads attend to the entire recording, while others focus on specific aspects or trials in the data. These results suggest that the fingerprints of each cell type are distributed across different time scales.

**FIGURE 3:**
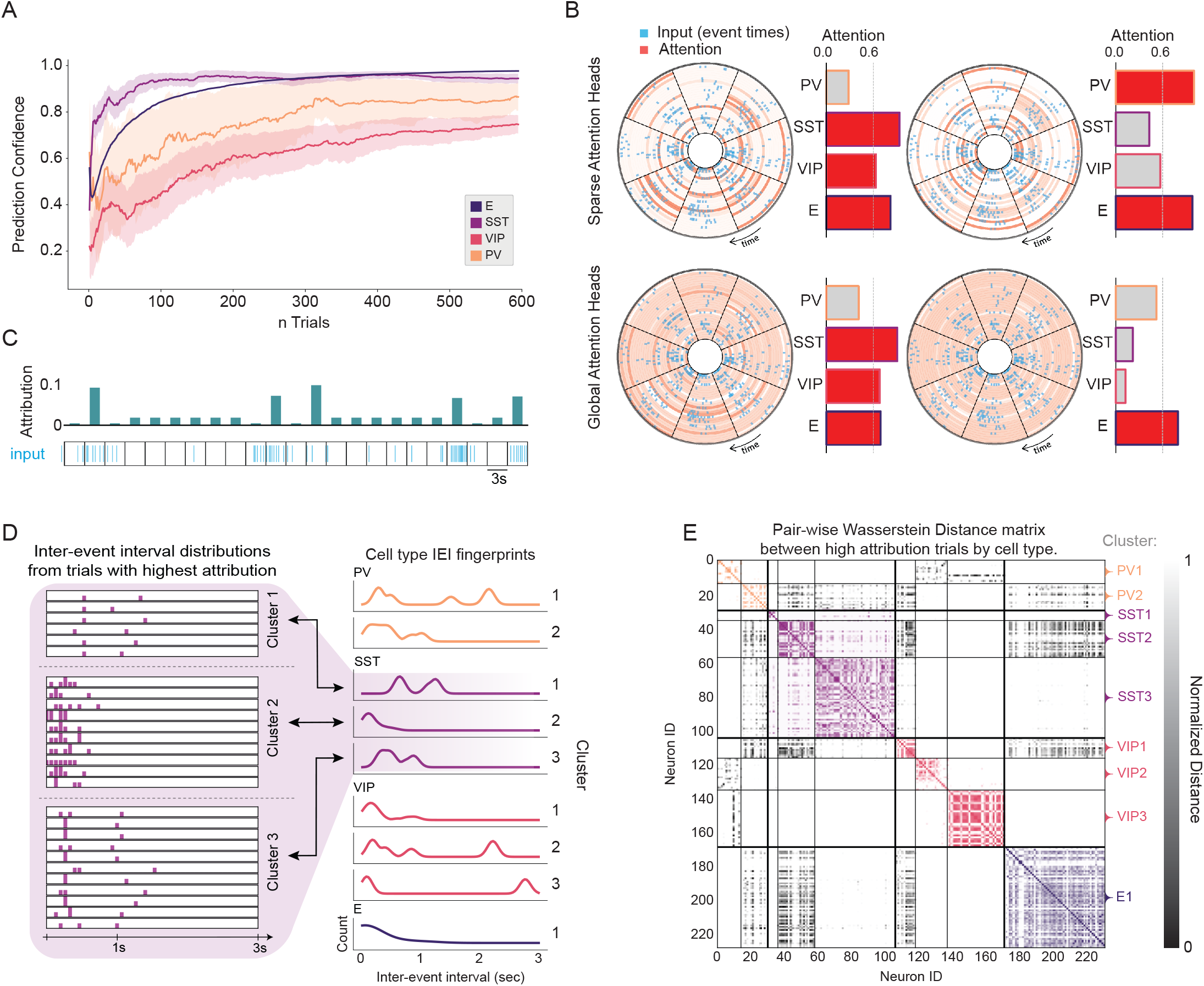
Interpretability of computational fingerprints learned by LOLCAT in calcium imaging data. A) Prediction confidence by cell class as a function of the number of observed time windows. B) Visualization of attention weights across the four heads in the attention block organized according to whether they attend to different time points in a global (bottom row) or sparse (top row) manner. A single neuron’s firing raster is displayed (blue), with each of the eight slices (black lines) denoting the presentation of a stimulus. Trials are further organized and stacked in concentric circles (of different radii). Bar plots show accuracy of decoding cell type from each head individually (rather than concatenating all four heads as in the full LOLCAT model). No one head individually decodes all cell types; the success of LOLCAT emerges from the combination of the different signatures. C) Integrated Gradients (IG) is applied to LOLCAT to reveal an attribution score (teal bars; top), corresponding to the relative importance of each trial for LOL-CAT’s prediction of the correct class. IEI raster (blue; bottom) corresponding to the attribution score shown above. D) IEI distributions for high attribution trials reveal consistent fingerprints of cell classes after clustering with the Wasserstein Distance. Representative trials from each of the three SST clusters is shown on the left with the Wasserstein barycenter (center of mass) displayed to the right. Excitatory neurons exhibit unimodal fingerprints while interneurons reveal bi-modal structure containing both fast and slower activity dynamics. E) The pairwise Wasserstein divergences between the distributions of trials with highest attribution, ordered by cell type and cluster. While all cell classes exhibit some similarity with other classes, PV appears to be the most distinct in terms of IEI fingerprints.

To examine the specific local features that contribute to different cell type classification, we applied the integrated gradients (IG) method (Sundararajan et al., 2017), a well known method for interpretability in deep neural networks; we applied IG to highlight salient trials that were useful for the network’s prediction of different cell types (Fig. 3C,D) and built an average fingerprint for neurons in each class (see Methods for details). This analysis revealed different properties distinct to each cell type. In particular, we found that inhibitory cell classes have multi-modal peaks in spiking and that salient trials for excitatory neurons tend to be unimodal and concentrated in shorter interevent intervals. K-means based clustering of IEI fingerprints revealed distinct subtypes of fingerprints within the three inhibitory classes but not excitatory (Fig. 3D). The local fingerprints associated with each cluster exhibited heterogeneous patterns of similarity across groups (Fig. 3E), further suggesting that additional global information is necessary to extract genetic class.

### Excitatory subtypes

The inhibitory classes considered here are genetically distinct and are stable over time (Scala et al., 2021). In other words, PV neurons do not express SST, and are unlikely to switch classes. In contrast, whether subtypes of excitatory neurons are distinct and stable is an important question under active investigation. The transcriptomic signatures that define excitatory classes may be better thought of as a gradient (Scala et al., 2021). For example, activity during development may drive the emergence of various cell types within cortical areas (Cheng et al., 2022; Tasic et al., 2018; Nowakowski et al., 2017). This raises questions of their separability by a computational fingerprint, let alone transcriptomics. We took advantage of the 10 excitatory Cre lines contained in the calcium imaging dataset to address this question, excluding two lines, EMX1 and SLC17a7, known for their widespread glutamatergic expression and lack of specificity (Gorski et al., 2002; Tasic et al., 2016; Daigle et al., 2018). We first trained LOLCAT to identify and separate excitatory neurons from inhibitory neurons (K=4, Fig. 2D). We then trained a LOLCAT with K=8 on the excitatory neurons to understand which, if any, excitatory subtypes produce a distinct computational fingerprint (Fig. 4A). Overall, we found that certain lines like Cux2, Ntsr, and Rbp4 could be discerned at or near over 50% accuracy (chance is 12.5%), while other lines were confused with a subset of other possible lines. Specifically, two subtypes, Rorb, and Scnn1a performed below 30%, and have been demonstrated to label cells more broadly than highly specific transcriptomic subclasses (Tasic et al., 2016; Matho et al., 2021), such as Cux2 (52%)(Gil-Sanz et al., 2015) and Ntsr (50%) (Harris et al., 2019). These data indicate that genetic specificity of excitatory cell types is correlated with the resolvability of cell types in the time series of neuronal activity.

**FIGURE 4:**
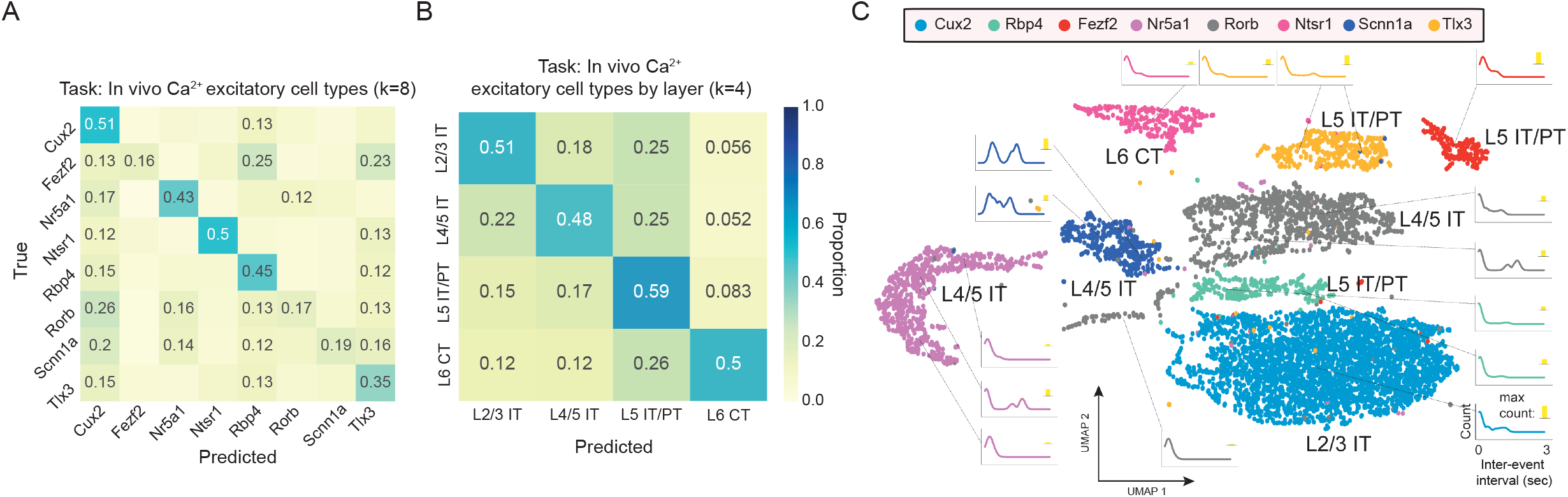
Breakdown of excitatory genetic subtypes within in vivo calcium imaging. A) Confusion matrix for LOLCAT applied to the time series of neuronal activity recorded in eight excitatory Cre lines. Annotation of performance at or below chance is deleted for visual clarity. B) Confusion matrix for LOLCAT applied to excitatory cell type when the 8 CRE lines are grouped by laminar position and recording depth (Layers 2/3, 4, 5/6). C) Uniform manifold approximation and projection (UMAP) embedding of all separable cells colored by their respective Cre lines. Layer/depth assignment and projection class are indicated by adjacent labels (intratelencephalic: IT; pyramidal tract: PT; corticothalamic: CT). In each class, we visualized interevent interval fingerprints for each cluster and show the Wasserstein barycenter of the IEI distributions of nearby cells. Inset yellow bars indicate the relative magnitude of the peak of the IEI fingerprint. Here, we see far more complex fingerprints for excitatory cells than in the 4 class condition, where excitatory types had relatively simple unimodal distributions.

As further distinctions within canonically defined cell types are still being recognized and genetic specificity of excitatory Cre lines is variable, we collapsed the excitatory predictions based upon both layer and projection class (Yao et al., 2021; Harris et al., 2019). This resulted in four excitatory types: Layer 2/3 IT neurons (Cux2), Layer 4/5 IT neurons (Nr5a1 - L4, Rorb - L4/5 IT, Scnn1a - L4/5 IT), Layer 5 IT/PT (Tlx3 - L5 IT, Rbp4 - L5 IT/PT), and Layer 6 CT neurons (Ntsr1) (Fig. 4B). In this context, LOLCAT achieved 52.1 +/-0.3% balanced accuracy. The highest accuracy (59.5%) was achieved for neurons in layer 5 IT/PT and the other classes reach roughly 50% accuracy, suggesting that excitatory cell types may be more easily differentiated based upon their layer and projection type than cell type.

We visualized the representations learned by the model with a supervised variant of uniform manifold approximation and projection (UMAP) that aims to keep points from the same class close in the embedded space (Fig. 4C)(McInnes et al., 2018). In this embedded space, we computed the Wasserstein barycenter (center of mass) of the top IEIs (see Figure 3) extracted for nearby groups of neurons. In contrast to our previous analysis of excitatory IEI fingerprints for K=4 when all excitatory cells are considered as a single group (Fig. 3D), these analyses revealed distinguishable and complex fingerprints within the excitatory cell class population. Of particular interest are the layer 4 Cre lines which exhibit multi-modal fingerprints when compared to the other excitatory classes.

### Generalizability of fingerprints across stimuli

We next sought to determine whether the computational fingerprints of cell type are specific to the stereotyped and constrained stimuli that comprise drifting gratings. Visual neuronal responses in one set of conditions are not predictive of responses in other conditions (de Vries et al., 2020), thus it is unclear if generalizable principles exist in neuronal spiking. To address this, we trained a LOLCAT to predict cell classes (K = 4; chance = 25%) in the calcium imaging dataset collected during presentation of drifting gratings. As shown in Fig. 2D, this model performed with high balanced accuracy when tested on withheld neurons also in the context of drifting gratings (Figs. 5A). The same model performed relatively poorly when tested on neuronal time series collected during the presentation of natural movies (mean balanced accuracy 48.5%), with a 61% range of class accuracies, exhibiting a tendency to overpredict excitatory neurons (Fig. 5B). However, when we evaluated the component attention heads individually, those that learned sparse rules (i.e., honed to specific trials and not global features) performed far better than the complete model, learning to classify three of the four classes with > 70% accuracy (Fig. 5B). To understand why global rules learned during drifting gratings might be deleterious to the performance of the same model tested during naturalistic movies, we examined the context-dependence of the simplest summary statistic of neuronal activity: rate. Compared to drifting gratings, event rates during naturalistic movies shifted markedly upwards in VIP and PV neurons, slightly upwards in excitatory neurons, and downwards in SST neurons.

**FIGURE 5:**
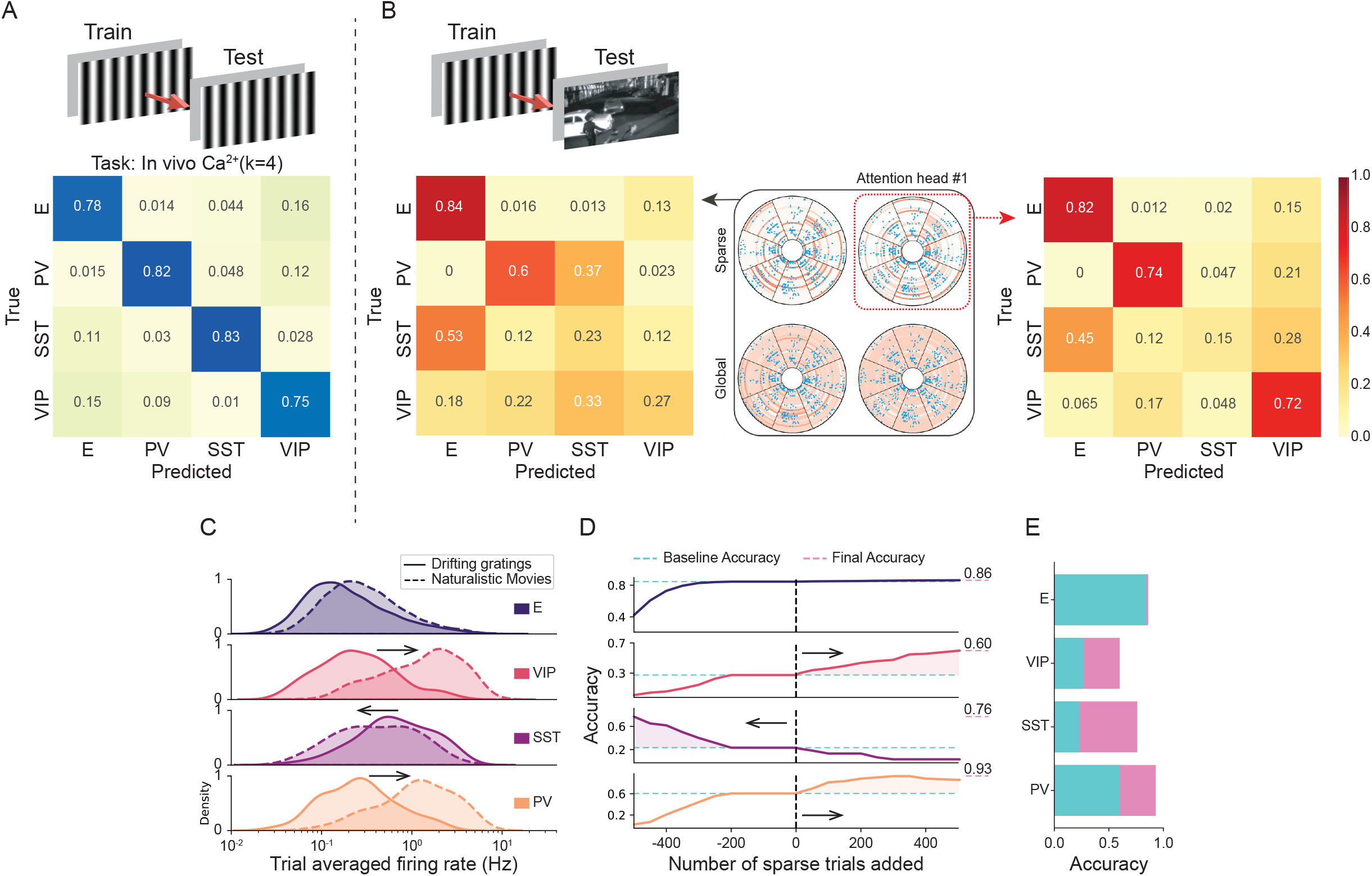
Generalizability of computational fingerprints across stimulus sets. A) The LOLCAT model is trained on neural activity recorded during the presentation of drifting gratings. Train/test stimuli are shown (top) and the corresponding confusion matrix for K = 4 cell classes. B) Models trained on drifting gratings and tested on natural movies (top). The total model performance (left confusion matrix), which includes locally and globally tuned attention heads (center), is outperformed by individual, locally tuned attention heads (right confusion matrix). C) Neuronal event rates, a global parameter, measured during gratings and naturalistic movies, exhibit cell class specific shifts in distribution. D) LOLCAT accuracy by cell class as a function of adding or removing sparse trials to account for global changes in event rates. E) LOLCAT accuracy by cell class prior to event rate correction (teal) and following correction (pink).

To test whether changes in the baseline rate of activity were sufficient to alter model performance, we systematically altered the number sparse trials within each cell-type when testing the drifting gratings-based model on neuronal activity acquired during natural movies (Fig. 5D). Specifically, in PV and VIP classes, firing rates were higher during natural movie presentation than in drifting grating presentation. We compensated for this by replacing activity from some dense (high-activity trials) with sparse (low-activity trials) prior to classification. For SST classes, sparse trial activity was replaced with dense trial activity instead. Compensating for the basal rate of each cell class drove significant increases in the generalizability of the LOLCAT model across stimuli in all but excitatory neurons, the only class well identified originally (Fig. 5E). SST neurons in particular exhibited a 50% improvement in prediction. Taking rate into consideration resulted in a mean accuracy of 78.8% when generalizing across stimulus sets. This was within 1% of the same model (trained on drifting gratings) tested on drifting gratings (79.6%; Fig. 2D,F; Fig. 5 E). Taken together, these results support the hypothesis that consistent transcriptomic class information is embedded in the time series of neuronal activity across diverse contexts.

## DISCUSSION

In this study, we asked whether a neuron in a complex cortical network embeds a fingerprint of its transcriptomic cell type into its activity. To test this, we examined the resolvability of different cell types using spike and event timing in 1) a bio-realistic model of visual cortex, 2) an open dataset comprising in vivo high density extracellular electrophysiological recordings and 3) an open, in vivo optophysiology dataset from mouse visual cortex. Taking advantage of the fact that many of the recorded neurons were labeled according to their transcriptomic cell type, we sought to answer a fundamental question: does cell type constrain a neuron’s spike timing in the intact circuit? By considering the largest dataset of cell type-labeled neuronal activity available, we demonstrate that standard metrics fail to meaningfully separate cell types, even when considered in a model with 230,000 neurons assigned to distinct types. However, by deploying a novel deep learning architecture that learns across time scales, we show that it is possible to decode subclasses of both inhibitory and excitatory neurons from the event times of individual neurons. In in vivo electrophysiological recordings containing labeled inhibitory classes (SST, PV, and VIP), we achieved a >50% accuracy for each class. In in vivo calcium imaging, we achieved >=75% accuracy for the three inhibitory and an excitatory class. Within the excitatory class, we find that specific excitatory types are variable in their distinguishability, and that considering layer and projection class may offer a more reliable means of classifying computational fingerprints in excitatory neurons. Our study reveals that a reliable stream of information about cell type is embedded within the time series of neuronal activity in the intact brain of behaving animals.

Linking transcriptionally defined neuronal types to their functional phenotypes in the intact brain is a widely recognized challenge (Scala et al., 2021). Traditionally, the cellular diversity of neurons was described in terms of physiological and anatomical features (Harris and Shepherd, 2015). Currently, neural taxonomies rely on either transcriptomic or morpho-electric parameters (Ecker et al., 2017; Mukamel and Ngai, 2019; Scala et al., 2021; Gouwens et al., 2020). Most often, neuronal cell type classification depends on molecular markers (Rudy et al., 2011; Kepecs and Fishell, 2014; Tremblay et al., 2016) or transcriptional profiles (Zeisel et al., 2015; Tasic et al., 2018; Zeisel et al., 2018). However, these approaches are either limited to a single cell type or provide limited insight into the functioning, intact circuit. In the intact circuit, a loose reflection of cell type is suggested by clustering neurons based on the shape of the extracellular waveform. Depending on species and brain regions, waveforms reveal two to eight groups (Lee et al., 2021; Henze et al., 2000; Jia et al., 2019; Trainito et al., 2019; Hengen et al., 2013; Bartho et al., 2004), but with the exception of putative parvalbumin-positive interneurons, the mapping of waveform onto transcriptomic class is unknown. As a result, parallel progress in transcriptomic classification of cells and technology for recording from large numbers of neurons has not yet emerged. Our results indicate that LOLCAT and other deep learning architectures (e.g. MLP) will enable the rapid identification of tens of thousands of simultaneously recorded neurons observed during behavior and cognition.

It is unclear that a neuron’s spike timing in vivo should be meaningfully and differentially informed by cell type. While in vitro and ex vivo results have identified physiological differences between cell types (Gouwens et al., 2020), these effects are not necessarily detectable in the context of the intact brain. One possibility is that, once wired to appropriate pre- and post-synaptic targets, a neuron’s genetic cell type does not systematically constrain spike timing. Even in this case, correlations across networks of cells could reveal the influence of cell type, as shown by Bugeon et al. (2022). Prior efforts to use spike timing to resolve even small numbers of excitatory cell types *in vivo* concluded that network driven variability in neuronal responses conceals the existence of subtypes (Crockett et al., 2015). Further, there is a long literature centered on the randomness of the activity of isocortical neurons (Ma et al., 2019; Shadlen and Newsome 1994, 1998); the spike times of isocortical neurons appear to be Poisson in nature, a feature that provides computational robustness (Van Vreeswijk & Sompolinsky, 1996; Ma et al., 2019) and appears incompatible with reliable, cell-type specific responses. However, our work may reconcile prior observations of chaotic dynamics in vivo and physiological differences in cell types ex vivo. Consideration of individual neuronal activity across multiple time scales (ms to >30 minutes, in this case) is sufficient to decode cell type well above chance in the context of a global Poisson distribution. This suggests that cell type meaningfully and consistently influences activity of a neuron, even in a chaotic, recurrent network faced with externally driven inputs. This observation raises the question of whether neurons take advantage of information about the identity of their inputs. Previous modeling suggests that this type of information may be utilized by sensory neurons (Gütig and Sompolinsky, 2006); further work will be necessary to clarify whether the brain utilizes information streams on longer time scales that carry cell identity information.

Intrinsic in the architecture of LOLCAT is an absence of assumptions about relevant time scales that might support a functional fingerprint of cell type. The success of LOLCAT implies that cell type information could manifest in the timing of neuronal activity in one of two ways. First, it is possible that most snippets of local activity (e.g., a trial) are not informative, but a select subset contain a unique fingerprint that is detected through our proposed attention mechanism. Second, it is possible that cell type influences the reliability of local activity and a more robust signature of cell type can be built when local activity is considered over longer time scales. In each of these cases, a single trial’s worth of activity will rarely be informative, and the summary statistics of activity could be entirely overlapping, yet cell type be readily extractable by LOLCAT. By visualizing the performance of the four attention heads in the architecture, it is clear that information about cell type is distributed according to *each* of these principles, and the balance between them differs depending on cell class. Intriguingly, this balance (summarized as the number of trials necessary to predict dell type) is itself an indicator of the K=4 classification of cell class. That SST interneurons require the fewest local observations suggests that these cells are highly consistent in their patterns of activity, and that these patterns are largely unique to the class. This parallels prior observations of the reliability of SST local activity (de Vries et al., 2020).

While the LOLCAT architecture is able to learn non-linear relationships that exist between event times and cell type labels, we developed interpretability methods to provide insight into local features (salient trials) as well global attributes (attention weights) used to classify different cell types. At the level of individual trials, interesting features of local IEI distributions are used by the model, i.e., whether the event arrival times exhibit unimodal or multimodal structure. In our K=4 comparison (where excitatory cells are considered as a class), the model highlights complex local structure for inhibitory classes and relies on a unimodal IEI distribution to identify E cells. However, when we break down the excitatory class into types and learn a new model to classify the 8 E Cre lines, more complex local features are discovered in Layer 4 neurons, and diverse responses are highlighted in our analysis. When analyzing more global features, certain classes are well predicted in a single attention head, while other classes are distributed across multiple attention heads. Interestingly, neurons can be predicted in all of the heads considered independently and in cases where other cells are poorly predicted (bottom right, Figure 3B); in contrast, PV neurons are best predicted in a single sparse head. SST neurons are predicted in both a sparse and dense head and can be easily predicted from a small number of trials (Figure 3A).

The generalizability of cell type fingerprints might be facilitated by deliberately limiting LOLCAT’s ability to contextualize neuronal activity. We use IEI distributions to represent neuronal activity (rather than an ordered sequence of spike times) and allow our multi-head attention module to pool representations agnostic to the temporal order of trials. One effect of this is that the model does not have access to the information needed to align spike trains across neurons on both short and long timescales (i.e. identify temporally correlated activity). Further, by passing each trial’s activity through an identical encoder, we limit the network’s ability to link patterns of activity to specific stimuli (i.e. tunings). These limitations in representation may serve to regularize the network, avoiding overfitting to specific contexts. Thus our results expand upon the connections of transcriptomic cell type to population correlations and neuronal tunings (Bugeon et al., 2022).

This work advances our understanding of cell type, revealing patterns in a neuron’s timeseries can reliably discriminate both inhibitory classes and excitatory cell types across diverse contexts. The ability to infer transcriptomic cell type from recorded neuronal activity opens up a wide range of possibilities. Capitalizing on the novel observation that transcriptomic types and classes drive computational differences, the interactions of many cell types can be estimated in the context of complex cognition and behavior. Insights could be integrated into the architecture of computational models of cortical processes or influence the design of experiments targeting individual or multiple cell lines. In addition, given the important and evolving connections between neuroscience and artificial intelligence (AI) (Richards et al., 2019), these results also raise the possibility that inclusion of different cell types as unique components in deep learning architectures will provide a computational benefit for AI.

## MATERIALS AND METHODS

### 1. Datasets

#### 1.1 Neuropixels datasets

Neural data from the Allen Brain Institute’s Visual Coding - Neuropixels dataset was retrieved as NeuroData Without Borders (NWB) files through AllenSDK. The default filter criteria for selecting quality units was applied. For recordings coinciding with all presentations of drifting gratings, high quality units whose average firing rate was at least 0.1 hz were included for analysis. This did not eliminate classes from the dataset. For drifting gratings, we retrieved recordings coinciding with all 3 second stimulus presentation trials. In the Brain Observatory Dataset, stimulus presentations varied in spatial orientation (0º, 45º, 90º, 135º, 180º, 225º, 270º, 315º, clockwise from 0º = right-to-left) and temporal frequency (1, 2, 4, 8, and 15 Hz), remained consistent in spatial frequency (0.04 cycles/degree) and contrast (80%) with 15 equivalent presentations of each particular stimulus. They were followed by 1 seconds of gray screen. In total, this amounted to 30 minutes of recording from each unit. In the Functional Connectivity Dataset, stimulus presentations varied in spatial orientation (0º, 45º, 90º, 135º) and 9 contrasts (0.01, 0.02, 0.04, 0.08, 0.13,0.2, 0.35, 0.6, 1.0), remained consistent in temporal frequency (2 Hz) and spatial frequency (0.04 cycles/degree) with 75 equivalent presentations of each particular stimulus. They were followed by 1 seconds of gray screen. In total, this amounted to 30 minutes of recording from each unit.

#### 1.2 Calcium Imaging datasets

Neural data from the Allen Brain Institute’s Visual Coding - 2-Photon Imaging dataset was retrieved as NeuroData Without Borders (NWB) files through AllenSDK. The default filter criteria for selecting quality units was applied. For recordings coinciding with all presentations of drifting gratings, high quality units with five or more events during at least one trial were included for analysis. This did not eliminate classes from the dataset. For drifting gratings, we retrieved recordings coinciding with all 3 second stimulus presentation trials. Stimulus presentations varied in spatial orientation (0º, 45º, 90º, 135º, 180º, 225º, 270º, 315º, clockwise from 0º = right-to-left) and temporal frequency (1, 2, 4, 8, and 15 Hz), remained consistent in spatial frequency (0.04 cycles/degree) and contrast (80%) with 15 equivalent presentations of each particular stimulus. They were followed by 1 seconds of gray screen. In total, this amounted to 30 minutes of recording from each unit. For naturalistic movies, we retrieved recordings coinciding with presentations of Natural Movie 3, a 120-second clip with no cuts from the movie “A Touch of Evil”. This was presented 10 times, amounting to 20 minutes of recording from each unit.

Separately for recordings coinciding with all presentations of drifting gratings or naturalistic movie, high quality units with at least 5 events in one or more subsequent 3-second intervals (i.e. trials for drifting gratings) were included for analysis. For some tasks, this eliminated classes from the dataset.

#### 1.3 Biorealistic model for generating synthetic spike trains (V1)

We have used the Allen Institute’s V1 model for simulations of mouse primary visual cortex (Billeh et al. 2020). The V1 simulations are done using Generalized leaky integrate-and-fire (GLIF) models, which have the same connectivity graph as the biophysical model (Billeh et al. 2020). This model has a core of radius 400 µm surrounded by leaky integrate and fire model neurons, totaling a radius of 845 µm. For more details please see Fig 1B of Billeh 2020. The V1 model contains 51,978 cells in the core and the whole network contains 230,924 cells with 85% excitatory and 15% inhibitory neurons. The V1 network contains 17 cell types represented by 111 unique GLIF neuron models. After filtering for neurons with firing rate exceeding 0.1 hz, layer 5 inhibitory Htr3a were very rare and therefore not included in classification tasks. This simulation consists of 100 presentations of drifting gratings to LGN with different orientations (0, 45, 90, 135, 180, 225, 270, and 315). Each of these 100 trials starts with 500 ms for grey screen followed by drifting gratings in an orientation angle for 2.5 s. The total duration of the simulation was 300 s.

### 2. LOLCAT

#### 2.1. Method

Our approach for cell type identification, LOLCAT, consists of three main components. We outline the architecture and rationale behind each of these components below.

##### (1) Local feature extractor

We observe the response of the neuron to a series of stimulus presentations, the length of each trial is 3 seconds, but the total number of trials slightly varies across the dataset. We first process each trial individually before aggregating the information globally: For each trial, the inter-event interval (IEI) distribution is computed using d log-spaced bins.

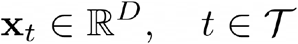

The resulting d-dimensional input vector is fed to the local feature extractor, which is a multi-layer perceptron (MLP) equipped with batch normalization layers and rectified linear units.

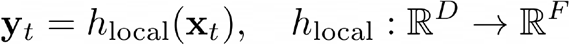

The same local feature extractor is shared across trials, as it is tasked to extract features that locally characterize the signature of a neuron’s activity. This design is directly influenced by the structure of our data (Battaglia et al. 2018). The reuse of the same feature extractor is known as weight sharing, and has the advantage of artificially augmenting the number of samples that the local feature extractor is trained on, as it observes one 3-second trial at a time, instead of observing the entire activity of the neuron at once. For the V1 k=16 task, the dimensions of the encoder was (128, 64, 64, 32) and for the k=4 task, the dimensions were (32, 16, 16, 16). For the neuropixels tasks, we perform a hyperparameter sweep on the validation set to determine the dimensions of several hyperparameters, and report the accuracy of the best validation hyperparameters on the test set.

##### (2) Multi-head attention module

We aggregate the extracted local features to produce a cell-level representation that describes its global distribution. To allow the network to seek out or attend to specific trials, we use an attention network (Bahdanau, et al. 2014; Li, et al. 2016) that generates an attention score for each trial, and then uses it to weigh the trial’s contribution to the final global feature vector.

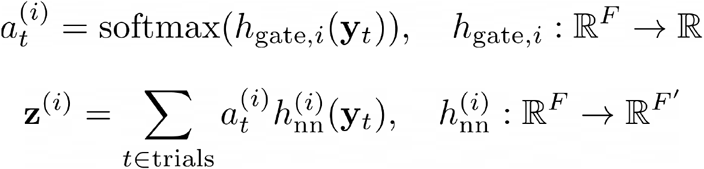

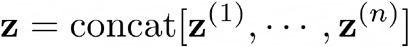

This is simply a weighted sum of the trial-level local features. The use of the softmax operator ensures that the attention scores sum up to 1. All these design choices enable us to apply our model to a cell with any number of observed trials, as this pooling operation does not depend on a fixed number of trials.

During training, the model can learn to attend to all trials equally in order to extract information about the average firing rate for example, or it can learn to be highly selective and identify trials that are relevant for distinguishing between the cell types, but the informational content of which would otherwise be drowned if all trails are considered equally. A single attention head assigns a single scalar to each trial, and thus can only learn a single rule on how to attend to the set of trials. To allow the model to learn different attention patterns, we use multiple attention heads (4 or 8) in parallel, each of which can learn to be selective to different activity patterns and aggregate statistics over different scales. All the pooled feature vectors are concatenated to produce a final global feature vector, which can simultaneously include features describing the average statistics of the neuron’s activity and other features encoding the presence of trials that are characteristic of a particular cell type (or group).

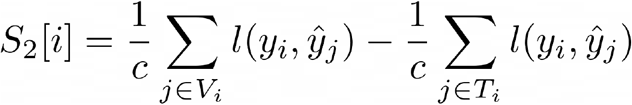

##### (3) Classifier

After concatenating all of the features from the different attention heads, we then pass this global representation to a final classification network, which uses the information aggregated at multiple scales, with different degrees of selectivity, to predict the cell type. We use an MLP with a single hidden layer.

#### 2.2. Training

We use He Initialization (He 2015) to initialize all of the model weights, except for the last layer of the classifier. To take into account the class imbalance in our datasets, we initialize the bias of the last layer of the classifier to be proportional to the class weight. This was found to speed up convergence.

We use the AdamW optimizer (Loshchilov and Hutter, 2017) with learning rate η and weight decay 10^{-5}. The learning rate is decayed once the training reaches the final epochs. We also use network dropout with a rate of 0.5.

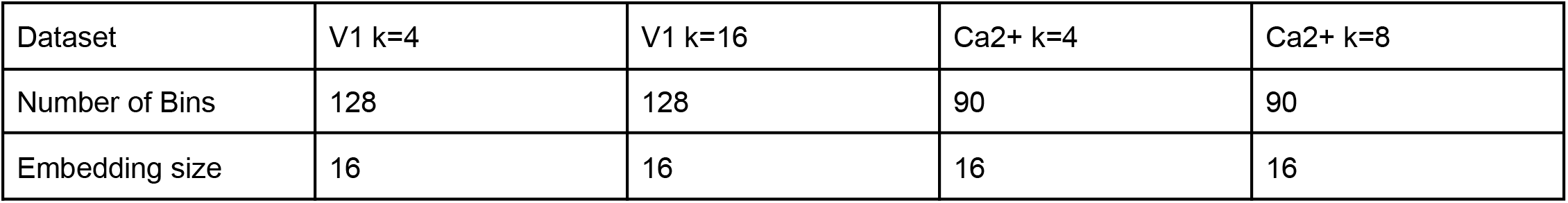
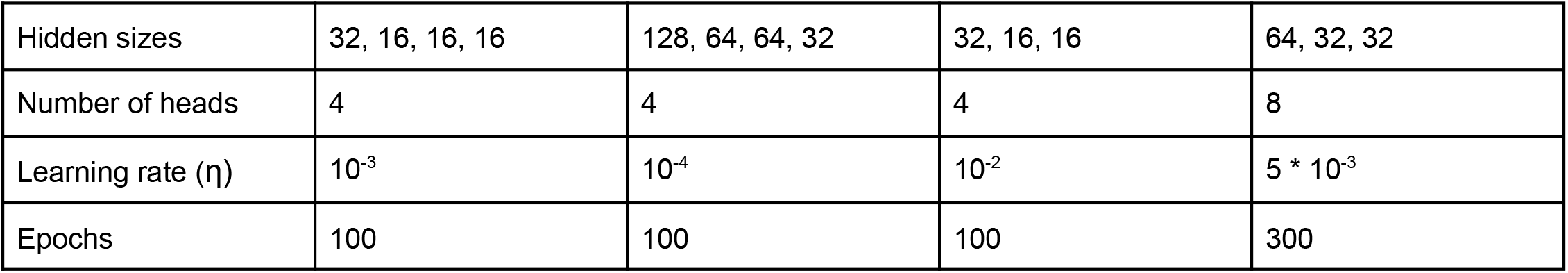

For the neuropixels tasks, we perform a hyperparameter sweep for each random seed. After an initial broad randomized search to determine the appropriate ranges for hyperparameter values on 5 splits, these smaller ranges were swept iteratively across each of the splits.

| Hyperparameter | Grid Values |
| --- | --- |
| Number of Histogram Bins | 64, 128 |
| Embedding size | 16 |
| Hidden sizes | (64, 32, 16), (64, 32, 16, 8), (128, 64, 32), (128, 64, 32, 16) |
| Number of heads | 4 |
| Learning rate | $10^{-2}$, $10^{-3}$ |
| Batch size | 64, 128 |
| Trial dropout | 0.3, 0.45, 0.6 |
| Network dropout | 0.5, 0.7 |
| Weight decay | $10^{-5}$ |
| Milestones | (50,100,200), (100,200) |
| Epochs | 300 |
| Sampler factor | 0.03, 0.1 |

#### 2.3. Data augmentation

Adaptive trial dropout is parametrized by dropout probability p. The probability of dropping a trial (removing it from the set of trials observed by LOLCAT), is proportional to p and inversely proportional to the neuron’s firing rate during that trial. In other words, if the trial is sparser than average, it will be more likely to be dropped out. This heuristic is inspired by similar heuristics designed for graph augmentations (Zhu et al. 2021).

#### 2.4. Dealing with class imbalance

Because of the class imbalance and variability across classes, we designed a strategy for reweighting different classes adaptively over training. To balance classes or decide the number of cells to sample from each class per batch, we calculate two scores: (i) a score that quantifies overfitting that might be taking place for a given class (ii) a score that quantifies how the model is performing on a given class relative to the overall dataset. At the end of each training epoch, we calculate the loss terms across both training and validation sets. First, we determine whether there is overfitting by calculating the difference between the average training and validation losses for a given class. If the difference is greater than 1-unit of standard deviation, calculated over all training loss terms, then the sampling factor for that class is increased for the following epoch (typically by division with 0.99). We also determine whether the model is over- or under-performing on a particular class by comparing the average per-class training loss to the globally averaged training loss, relative to the standard deviation. If the score is greater than a fixed threshold (0.1), then the sampling factor for that class is decreased (typically by division with 1.01) and otherwise increased. Initially, sampling factors for each class are set to balance the classes (majority class count/ class count).

More formally, the overfitting score can be expressed as follows:

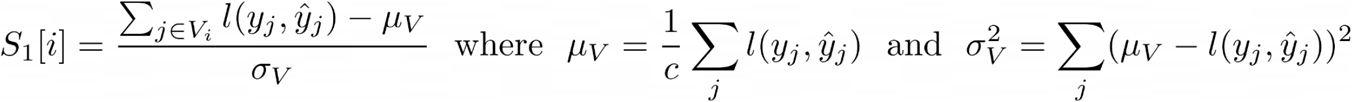

where the first term measures the average loss in classification prediction for all samples in the ith classes validation set and the second term measures the average loss in classification prediction for all samples in the ith classes training set. Thus, this score provides a gap in loss between the validation and train: when the model is overfit this score will be low.

The other measure we compute is how well a given class is predicted relative to cells in other classes. We compute this as follows:

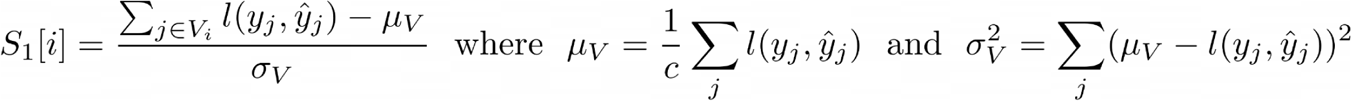

The first term in the score is the average loss on the validation set (as in our overfitting measure) and the mean and variance are computed over the entire validation set.

### 3. Baseline models

#### 3.1. Logistic regression model

For each neuron within a dataset, the mean firing rate (FR) and coefficient of variation (CV) were calculated. Using scipy.stats a gamma distribution was fit to the list of ISIs of the neuron and the alpha and beta parameters were extracted. For the spiking datasets (V1 model and Neuropixels), the distribution of ISIs below 500ms was used. For an average unit’s distribution, this yields roughly equivalent mean and standard deviation, which supports prediction of classes based on CV, alpha or beta. For the 2P-imaging datasets, the distribution of IEIs below 3s (the timespan of a trial was used). These basic statistics were used individually and collectively to attempt to predict class for each task. A logistic regression with penalized L2-Norm and 1E-4 tolerant stopping criterion was employed using sci-kit learn (Pedregosa et al., 2011).

#### 3.2. Multilayer Perceptron (MLP) model

For each neuron within a dataset, a binned histogram of the ISI distribution during blocks of drifting gratings presentation was used as features for an MLP implemented in scikit-learn (Pedregosa et al., 2011). For the spiking datasets (V1 model and Neuropixels) ISIs less than 500 ms were binned in 100 5 ms increments. For the 2P-imaging datasets, IEIs less than 3s were binned in 300 10 ms increments. In each dataset, a Bayesian hyperparameter optimization (implemented with the HyperOpt python package) was used to select network dimensions, regularization, learning rate, and class-balancing (using RandomOverSampling and/or RandomUnderSampling from the imbalanced-learn python package) to maximize balanced accuracy on the validation set (Bergstra et al., 2013; Lemaître et al., 2017). For several tasks, optimized class-balancing was critical to ensure minority classes were predicted. For all datasets, a single layer architecture of varying widths appears optimal.

### 4. Integrated Gradients and Clustering Approach for Finding Fingerprints

#### 4.1. Integrated gradients

The integrated gradients algorithm aims to explain the model’s predictions in terms of its input features, by identifying the features that were important in making the prediction possible (Sundararajan et al., 2017). We use a slightly altered version of the algorithm: we use it to explain the relationship between the model’s outputs and the trial-level features. In other words, we identify the trials that were critical in the prediction of the cell type. The algorithm assigns an attribution score to each trial. A trial with high attribution means that the neuron exhibited an activity pattern that skewed the model towards correctly predicting a specific cell type. This means that if this same pattern were to be observed, it is highly likely that the neuron is from the corresponding cell type.

#### 4.2. Clustering IEI distributions

For each cell type, we collect the inter-event interval distributions of the trials with the highest attribution across cells. Using the Wasserstein-2 distance as a measure of similarity between two distributions (Cuturi 2013), we perform clustering and then compute the Wasserstein barycenter of each cluster. The Wasserstein barycenter is essentially a different version of the centroid or average, where instead of taking a standard coordinate-wise average, the barycenter is the center-of-mass in the Wasserstein distance space and is thus the distribution where the Wasserstein-2 distance between all samples and the barycenter is minimized. We use this measure due to the sparsity of IEIs and small variability between different salient trials that when averaged together produces an overly smoothed version that doesn’t convey the shared structure between IEIs in a cluster as easily. Note that this analysis only focuses on the highly local information used by LOLCAT; the prediction is made after considering different pieces of information, aggregated across different scales.

## ACKNOWLEDGEMENTS

This project was supported by NIH BRAIN Initiative awards 1R01NS118442 (KBH) and 1R01EB029852 (ELD, KBH), as well as NSF award IIS-2039741 (ELD), and generous gifts from the Alfred Sloan Foundation (ELD), the McKnight Foundation (ELD), and the CIFAR Azrieli Global Scholars Program (ELD).

## AUTHOR CONTRIBUTIONS

Conceptualization, KBH; Methodology, KBH, ELD, MA, AS, KBN; Investigation, AS, MA, LMV, DP, and SE; Writing – Original Draft, KBH and ELD; Writing – Review & Editing, KBH, ELD, TN, and AS; Funding Acquisition, KBH and ELD; Resources, KBH and ELD; Supervision, KBH and ELD

